# Telomeric antisense oligonucleotides reduce premature aging phenotypes in telomerase mutant zebrafish

**DOI:** 10.1101/2025.05.23.655694

**Authors:** Giulia Allavena, Francesca Rossiello, Aurora Irene Idilli, Marina Mione, Fabrizio d’Adda di Fagagna, Miguel Godinho Ferreira, Bruno Lopes-Bastos

## Abstract

Telomerase activity is restricted in somatic cells, resulting in progressive telomere shortening. Telomere erosion eventually activates the DNA damage response (DDR), inducing cell-cycle arrest and cellular senescence or apoptosis. We previously reported that telomere dysfunction induces the transcription of telomeric non-coding RNAs (tncRNAs) which are critical mediators of DDR activation. Blocking tncRNAs with telomeric antisense oligonucleotides (tASOs) suppresses in vivo DDR signaling and its downstream effects. Here, we show that tASO-mediated inhibition of telomeric DDR in second-generation *tert−/−* zebrafish embryos with critically short telomeres leads to improved developmental outcomes and rescues premature aging phenotypes, including enhanced survival. Notably, a single tASO treatment administered at the one-cell stage of first-generation *tert*−/− embryos leads to enhanced fertility observed in 6-month-old adults. Overall, these findings demonstrate that tASO-based inhibition of telomeric DDR is sufficient to effectively rescue premature aging phenotypes in zebrafish.

---

Telomeres are the protective structures at the ends of chromosomes, composed in vertebrates of (TTAGGG)n DNA repeats and bound by shelterin complex proteins, which prevent recognition as DNA double-strand breaks (DSBs)^1^. When critically short, this protection at chromosome ends fails, activating the DNA damage response (DDR)^2^ and triggering cell-cycle arrest, cellular senescence, or apoptosis, all processes which contribute to tissue dysfunction, disease, and aging ^2–5^. Telomerase, a reverse transcriptase, counteracts shortening by elongating telomeres^6^, but its expression is limited in human somatic cells, leading to telomere loss with age. Similarly, telomerase mutant (tert−/−) zebrafish, lacking the telomerase protein component, display accelerated telomere shortening, premature aging phenotypes and reduced lifespan^3–5^. Thus, as in humans^7^, telomere length and integrity controls normal zebrafish aging ^4,5^, making it a valuable premature aging model. Interestingly, inhibiting DDR through tp53 mutation in tert−/− zebrafish rescues several telomere-associated defects^8,9^, suggesting DDR inhibition as a potential therapeutic strategy to extend healthspan. However, since p53 responds to various stressors, its inhibition may increase mutation rates and cancer risk, underscoring the need for more specific telomere DDR inhibition.

We previously reported that telomere dysfunction induces telomeric non-coding RNAs (tncRNAs), required for DDR activation at damaged telomeres^10,11^. Targeting these RNAs with telomere-targeted antisense oligonucleotides (tASOs) blocks DDR^10^, improves disease phenotypes, and extend survival in a progeria mouse model^12^. Here, we show that tASO-mediated telomeric DDR inhibition in tert−/− zebrafish improves development and rescues premature aging phenotypes, including fertility and lifespan with long-lasting effects. These findings demonstrate the efficacy of tASOs and support them as tools for research studies in telomere biology and aging and highlight them as potential therapeutic strategy for aging, accelerated-aging syndromes and age-related diseases.

## Results and Discussion

To study the physiological effects of persistent telomeric DDR caused by short telomeres, we used second-generation (G2) *tert−/−* zebrafish, which develop earlier and more severe phenotypes than first-generation fish (G1), making it an ideal model to test strategies to rescue short telomere phenotypes. G2 tert−/− fish exhibit much shorter telomeres than WT siblings (Fig.1A), along with increased inflammation and senescence^13,14^. In late-generation telomerase knockout mice, telomere shortening activates a p53-dependent DDR^15^. Similarly, we observed increased γH2Ax and p53 activation in G2 tert−/− larvae (Fig. 1B), demonstrating ATM-mediated DDR. We previously showed that short or damaged telomeres trigger a telomeric DDR through tncRNAs expression, which tASOs can supress^10,12^. Based on this, we tested whether tASOs could mitigate developmental effects of short telomeres in G2 tert−/− zebrafish.

**Figure 1:**
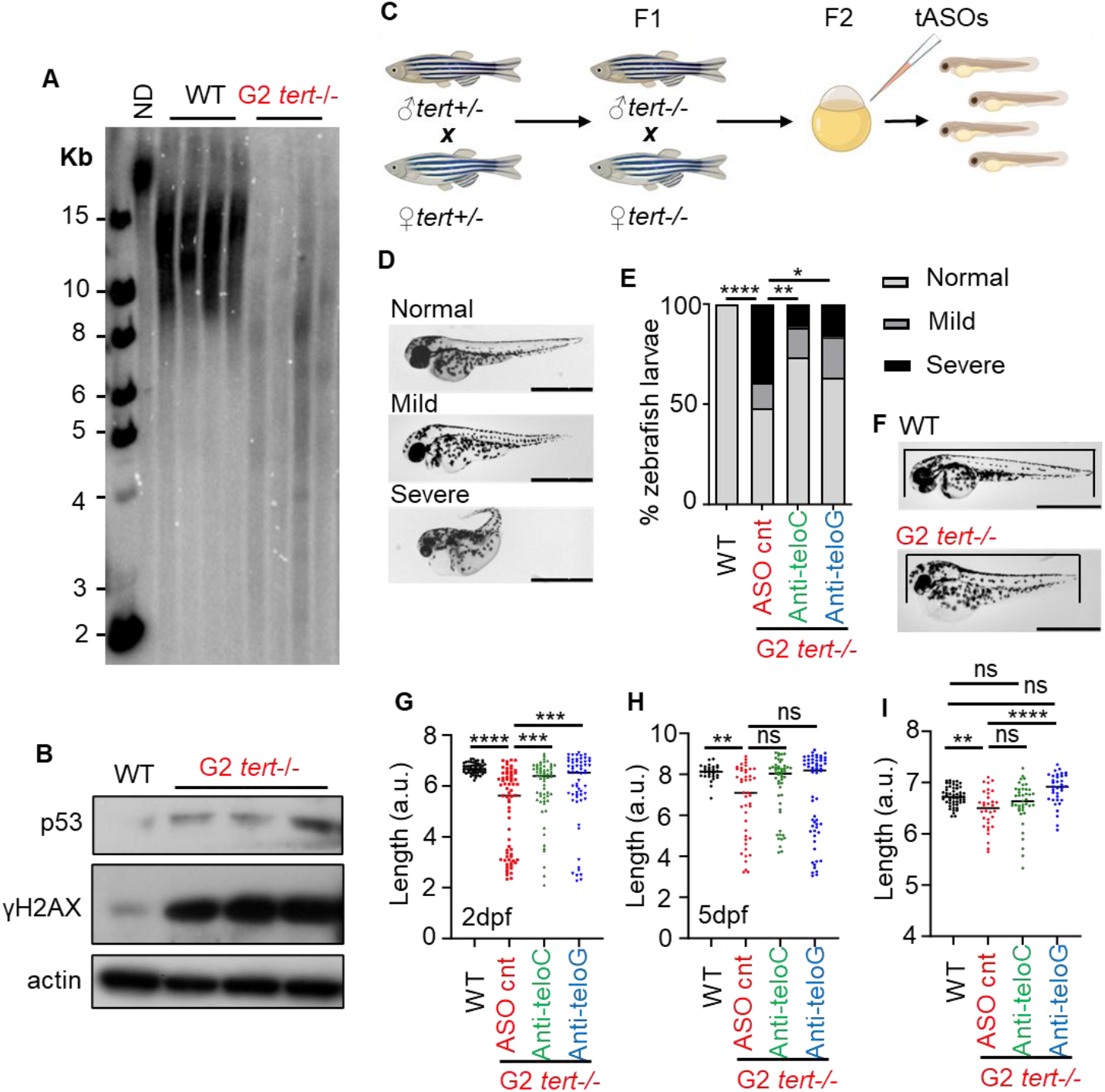
tASO treatment of second-generation tert−/− zebrafish reduces developmental defects. A- Telomere restriction fragment (TRF) analysis by Southern blotting of 5dpf WT and G2 *tert−/−* larvae, (ND: not digested DNA). B- Western blot of 5dpf WT and G2 *tert−/−* larvae. C- Experimental design scheme. D- Representative images and E- quantification of the phenotype in 2dpf WT (n=52) and G2 *tert−/−* larvae injected with ASO cnt (n=84), anti-teloC (n=60) or anti-teloG (n=60); Z-test. F- Representative images and G, H- quantification of larvae size in 2- and 5dpf WT (n=81) and G2 tert−/− larvae injected with ASO cnt (n=68), anti-teloC (n=60) or anti-teloG (n=60). I- Quantification of larvae size in 2-dpf WT (n=52) and in normal phenotype G2 tert−/− larvae injected with ASO cnt (n=32), anti-teloC (n=41) or anti-teloG (n=35). a.u. – arbitrary units; Line represents the mean and each dot represents one individual larva. One-way ANOVA. *p<0.05; **p<0.01; ***p<0,001, ****p<0,0001. Scale bars=1mm.

We injected tASOs complementary to either tncRNAs strand (anti-teloG complementary to (TTAGGG)n and anti-teloC complementary to (CCCTAA)n), or a control ASO, into one-cell-stage G2 *tert*−/− embryos (Fig. 1C). As expected, most control-treated G2 tert−/− larvae exhibited mild to severe malformations at 2 days post-fertilization (dpf), while WT larvae showed none (Fig. 1D-E). Notably, treatment with either tASO significantly reduced the percentage of *G2 tert−/−* larvae with severe phenotypes (Fig. 1E). Control-treated G2 tert−/− larvae were also significantly shorter than WT larvae at 2 dpf (Fig. 1F-G). Since larval length is a proxy for developmental stage, this suggests that short telomeres delay development. tASO treatment rescued larval length (Fig. 1G), and this trend persisted to 5 dpf (Fig. 1H). Although shorter size may result from high malformations rates, even excluding malformed larvae, G2 tert−/− larvae remained significant shorter (Fig. 2I), which tASOs fully restored to WT levels.

**Figure 2:**
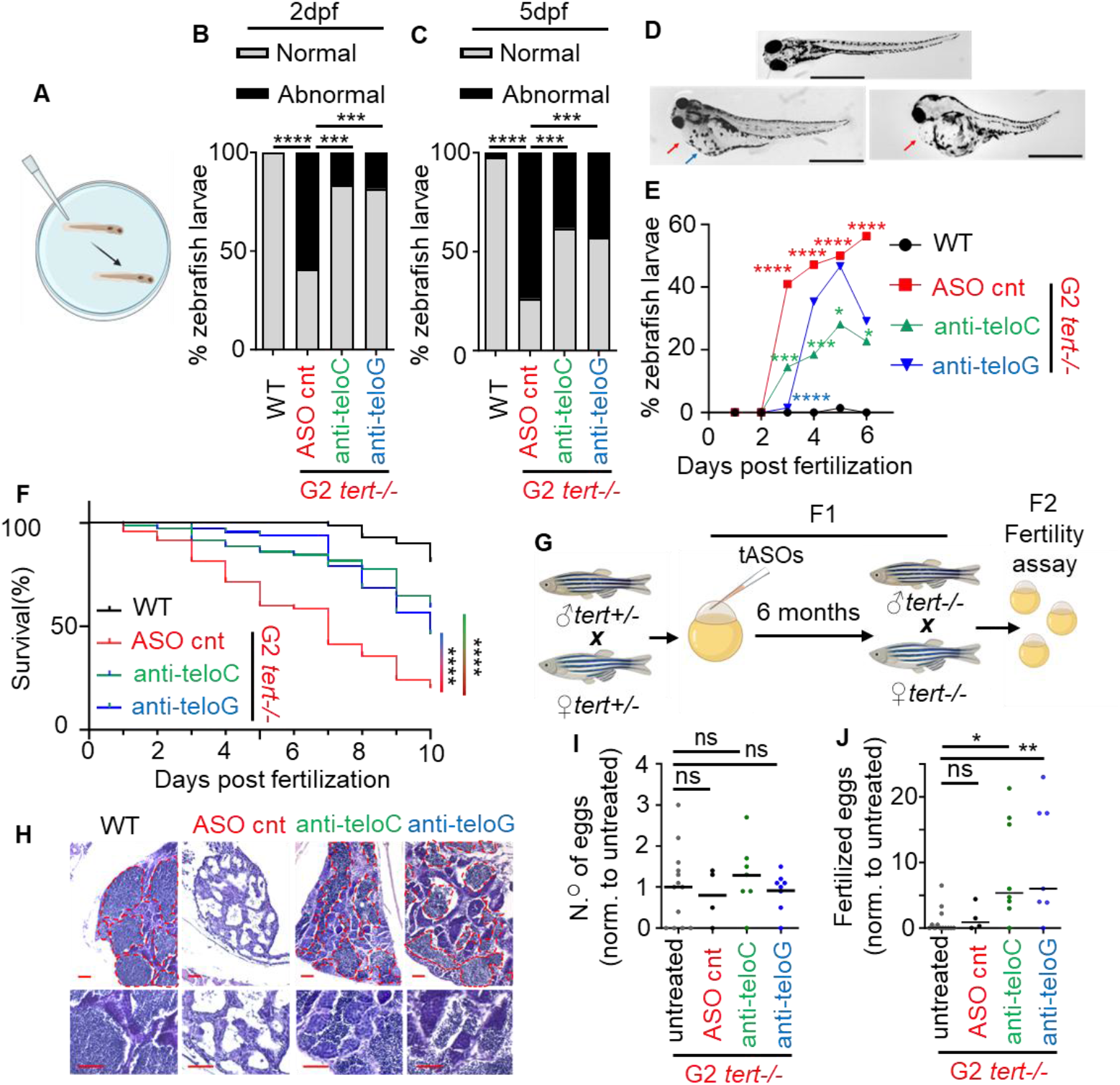
tASO treatment partial rescues detrimental phenotypes in tert−/− zebrafish with lasting effects. A- Schematic representation of behavior assay. B-C- Percentage of larvae displaying a normal or abnormal behavior to touch in 2 and 5dpf WT (n=55) and G2 tert−/− larvae injected with ASO cnt (n=32), anti-teloC (n=42) or anti-teloG (n=38). Z-test. D-Representative images and E- quantification of the percentage of larvae displaying edema in WT (n≥62) and G2 tert−/− larvae injected with either ASO cnt (n≥35), anti-teloC (n≥48) or anti-teloG (n≥42). Red arrow: pericardiac edema, blue arrow: yolk sac edema; scale bars=1mm. Z-test. F- Kaplan-Meier survival curve of WT (n=69) and G2 tert−/− larvae injected with either ASO cnt (n=56), anti-teloC (n=71) or anti-teloG (n=67). Log-rank (Mantel-Cox) test. G- Experimental design scheme. H- H&E representative images of testes of 6 months old fish. Red dashed lines outline mature sperm area. Scale bars=50µm. Quantification of I- total eggs and J- fertilized eggs laid by 6 months old G1 *tert−/−* zebrafish injected with tASOs at the one-cell embryo stage. WT (n=13), G2 tert−/− ASO cnt (n=4), G2 tert−/− anti-teloC (n=7), G2 tert−/− anti-teloG (n=8); One-way ANOVA. Line represents mean and each dot represents an individual cross. *p<0,05; **<0.01; ***p<0,001; **** p<0,0001.

These results demonstrate that short telomere-induced DDR impairs development and growth in G2 tert−/− larvae, and that tASO-mediated DDR inhibition can rescue these defects. This aligns with the high apoptosis levels reported in G2 tert−/− larvae^9^ and our previous work showing that tASOs decrease apoptosis and increases cell proliferation^12^, which can promote normal embryonic development.

Since our model is embryonic, distinguishing developmental defects from those linked to adult pathologies is challenging. We therefore assessed additional phenotypes likely linked to senescence and inflammation, previously observed in these embryos^13^. G2 tert*−/−* larvae showed reduced responsiveness to physical stimuli (Fig. 2 A-C), developed edemas (Fig. 2D-E) and had significantly shorter lifespan (Fig. 2F). Using a well-established behavioral assay^16^ (Fig. 2A), all WT larvae responded to touch with a normal escape reflex, while most G2 *tert−/−* larvae either failed to move or showed a swirling motion (abnormal behavior; Fig. 2B–C). Blocking telomeric DDR with tASOs increased the percentage of G2 *tert−/−* larvae with normal responses. Although we show that telomeric DDR is responsible for the abnormal behavior, the physiological causes remain unclear, we previously observed muscle fiber degeneration in G1 *tert−/−* zebrafish^4^, and studies in *tert* KO human motor neurons show impaired neurogenesis and age-related changes^17^, thus the swirling motion suggests a central nervous system dysfunction affecting motor coordination.

Another G2 tert−/− phenotype is the formation of edemas. Indeed over 50% of G2 tert−/− larvae developed pericardial and yolk sac edema after 2dpf (Fig. 2D-E), sometimes affecting also other body parts. Treatment with tASOs reduced edema (Fig. 2E), with anti-teloC having longer-lasting effect.

The most striking phenotype of G2 tert−/− larvae is increased lethality (Fig. 2F). Previous studies^3,14^, showed that most G2 tert−/− larvae die before two weeks. Treatment with either tASO significantly improved survival (Fig. 2F). At 10dpf, anti-teloG increased survival by 1.3-fold and anti-teloC by nearly 2-fold. This suggests telomeric DDR inhibition by tASOs not only improves individual phenotypes, such as length and malformations, but also enhances overall health allowing for increased survival.

To assess long-term effects, we evaluated fertility, which naturally declines with age in zebrafish and other mammals. In both murine and zebrafish *tert−/−* models, male fertility loss occurs earlier than in WT^3,5,18^. Since G2 *tert−/−* die before reaching sexual maturity, we used first-generation G1 tert−/− zebrafish injected with tASOs at one-cell stage (Fig. 2G). As expected, we observed testes atrophy in 6 months old fish, which was rescued by tASO treatment, assessed by an increase in sperm mature area (Fig. 2H). We also observed that, while in six-month-old G1 *tert−/−* fish tASOs treatments did not impact the number of eggs laid (Fig. 2I), they exhibited a fivefold increase in fertilized eggs compared to controls (Fig. 2J), demonstrating tASOs’ impact also on fertility months after treatment.

In this vertebrate model of accelerated aging, treatment with two distinct tASOs significantly reduced developmental defects, edema, behavioral abnormalities, and improved overall survival to nearly WT levels. Notably, a single tASO treatment had lasting effects, demonstrated by restored fertility six months post-injection. These benefits likely extend beyond apoptosis suppression. We previously showed *tert−/−* zebrafish exhibit increased senescence and inflammation^13^, which were reduced by DDR inhibition via p53 suppression^4,8^. In a progeria mouse model, tASOs also decreased both markers^12^, possibly contributing to the physiological rescue observed in tASOs-treated telomerase deficient zebrafish.

Overall, our results show that telomere-specific DDR inhibition can effectively counteract short telomere–associated phenotypes in zebrafish, supporting them as tools for research studies in telomere biology and aging in this model and as potential therapeutic agents for aging, accelerated-aging syndromes and age-related conditions.

## Material and Methods

One-cell-stage zebrafish embryos from an incross of either tert+/-, tert−/− or WT, were microinjected with 1 nl of 0.1 µg/µl tASO. Different assays were performed at the timepoints referred in the manuscript. Material and protocols are detailed in **SI Appendix**.

## Acknowledgements

This work was supported by Université Côte d’Azur—Académie 4 (Installation Grant: Action 2—2019), Agence Nationale de la Recherche ANR-21-CE14-0054 and La Ligue Contre le Cancer Equipe Labellisée 2024, France. B.L-B. was supported by a French FRM postdoctoral fellowship (SPF201809007006). F.d.A.d.F.’s laboratory is supported by: ERC Advanced Grant (TELORNAGING - 835103); ERC POC (TELOVACCINE - 101113229); AIRC-IG (30471); AIRC-IG (21762); AIRC 5×1000 (21091); Telethon (GMR23T2007); PRIN (2022R7LH5T); Next Generation EU, in the context of the National Recovery and Resilience Plan, Investment PE8 Project Age-It and Investment CN3 National Center for Gene Therapy and Drugs based on RNA Technology; FOE Virus-Memory (FOE 2020, FOE 2021). A.I. was supported by Fondazione Veronesi (postdoctoral fellowship - 2018)

## Supporting Appendix SI

### Extended Materials and Methods

#### Ethics statement

All animal experiments have been approved in accordance with national animal welfare guidelines in Portugal by the Ethics Committee of the Instituto Gulbenkian de Ciência and approved by the competent Portuguese authority (Direcção Geral de Alimentação e Veterinária; approval no. 0421/000/000/2015), in France by the Animal Care Committee of the Institute for Research on Cancer and Aging, Nice, the regional (CIEPAL Côte d’Azur no. 697) and national (French Ministry of Research no.27673-2020092817202619) authorities and in Italy according to D.Lgs. 26/2014 by authorization 148/2018-PR (Italian Ministry of Health) to M. C. Mione.

#### Zebrafish maintenance

Zebrafish were maintained in accordance with institutional and national animal care guidelines. The telomerase mutant zebrafish line *tert*^+/hu3430^ (referred to as *tert*−/−) was incrossed to generate G1 *tert*−/− individuals, which were then incrossed again to produce G2 *tert*−/− progeny. The *tert*+/− stock line was maintained by outcrossing to WT fish to prevent haploinsufficiency effects in the offspring. Noteworthy, that in vivo experiments were carried out by two independent laboratories (Mione’s and Ferreira’s) with consistent results.

#### tASOs injections

One-cell stage zebrafish embryos were microinjected with 1.4 nL of tASOs at a concentration of 0.1 µg/µL. Injected embryos were then incubated in E3 medium at 28 °C. tASO sequences were as follows (5−3′ orientation):

ASO control TTATCCGCTCACAATTCCACAT

anti-teloG ASO CCCTAACCCTAACCCTAACCC

anti-teloC ASO GGGTTAGGGTTAGGGTTAGGG

#### Telomere restriction fragment (TRF) analysis by Southern blot

A pool of 10-20 five dpf zebrafish larvae were sacrificed in 1g/L of MS-222 (Sigma Aldrich) and were lysed at 50^°C^ overnight in lysis buffer (Fermentas #K0512) supplemented with 1mg/ml Proteinase K (Sigma Aldrich) and RNase A (1:100 dilution, Sigma Aldrich). Genomic DNA (gDNA) was extracted using equilibrated phenol-chloroform (Sigma-Aldrich) and chloroform-isoamyl alcohol extraction (Sigma-Aldrich). The same amount of genomic DNA was digested with RSAI and HINFI enzymes (NEB) for 12 h at 37 ^°C^. After digestion, samples were loaded on a 0.6% agarose gel, in 0.5% TBE buffer, and run on an electrophoresis apparatus (110V for 15h, Bio-Rad). Gels were then processed for Southern blotting using a 1.6 kb telomere probe, (TTAGGG)n, labeled with [alpha-32P]-dCTP.

#### Western blot

A pool of 10-20 five dpf zebrafish larvae were sacrificed in 1g/L of MS-222 (Sigma Aldrich) and homogenized in RIPA buffer (sodium chloride 150 mM; Triton-X-100 1%, sodium deoxycholate 0.5%, SDS 0.1%, Tris 50 mM, pH=8.0), supplemented with protease and phosphatase inhibitor cocktail (Roche diagnostics) with a motor pestle on ice. Homogenized larvae were incubated for 30 minutes on ice and centrifuged at 4^°C^, 13 000 rpm for 10 minutes. 50 µg of protein/sample were loaded into a 10% SDS-PAGE gel and transferred to a Nitrocellulose membrane (BioRad #1620097). Membrane was first blocked with 5% skimmed milk, incubated with primary antibody (p53: 1:1000, Anaspec, 55342; γH2Ax: 1:1000, GeneTex, GTX127342; actin: 1:1000, Sigma-Aldrich, A2066) overnight at 4^°C^, and followed by secondary antibody (anti-rabbit, 1:10 000, Santa Cruz Biotechnology, sc-2357) incubation at room temperature for 2 hours. Chemiluminescence detection was performed with an ECL KIT (Amersham).

#### Behavior assay

A single larva was placed in the center of a Petri dish containing E3 medium. After one minute of acclimation, the tail was gently stimulated with a pipette tip, and the larva’s response was recorded. Larvae that moved forward in response to the stimulus were classified as exhibiting normal behavior, while those that failed to move or moved in a swirling pattern were classified as having abnormal behavior. Larvae displaying a severe phenotype were excluded from this assay, as they were unable to move independently.

#### Phenotype and length assessment and analysis

Larvae were raised in Petri dishes (maximum of 50 larvae per dish) in an incubator maintained at 28 °C with E3 medium, which was refreshed every two days. At 2 and 5 days post-fertilization (dpf), larvae were anesthetized with MS-222 (Sigma-Aldrich) and imaged using a Leica stereomicroscope. Larvae were scored based on phenotype severity as normal, mild, or severe. Larval length was measured using ImageJ.

#### Survival and edema assessment

Larvae were raised in Petri dishes (maximum of 50 larvae per dish) in E3 medium at 28 °C. The medium was changed every two days. Larvae were monitored daily, and the presence of edema and mortality was recorded.

#### Fertility assay

To assess fertility, individual breeding pairs (6 months of age) were housed overnight in separate external breeding tanks. The following morning, pairs were allowed to spawn and lay eggs. Eggs were then collected, and fertility was assessed between 4 and 6 hours post-fertilization.

#### Statistical analysis

Statistical analyses and graph generation were performed using GraphPad Prism version 10.1.0 (316). One-way ANOVA was used to compare the means of three or more experimental groups with normally distributed data. Z test was used to compare two or more groups of discrete data. The log-rank (Mantel–Cox) test was used to compare survival curves. The specific statistical test used for each graph, along with other statistical details, is described in the corresponding figure legend. A *p* < 0.05 was considered statistically significant throughout the study.

